# A synthetic biochemical device for sensing microgravity

**DOI:** 10.1101/2020.01.26.920629

**Authors:** Sayak Mukhopadhyay, Sangram Bagh

## Abstract

Biological solutions to human space travel must consider microgravity as an important component, which is unknown by the biochemical worlds on the Earth. Thus, one of the fundamental challenges of space biotechnology is to create engineered biochemical systems to integrate microgravity as a signal within molecular and cellular processes. Here we created the first molecular or biochemical microgravity sensor by creating a synthetic-small-regulatory-RNA based molecular network in *E.coli*, which sensed microgravity and responded by altering the expression of a target protein. We demonstrated that the design was universal, could work potentially with any promoter and against any target gene. This device was applied to target cell division process and rescue the deformed cell shape by applying microgravity. The work showed for the first time, a way to integrate microgravity as physical signals within biochemical process of a living cell in a human designed way and thus, opens a new direction in space biotechnology, space chemistry and space technology.

## INTRODUCTION

Rapid advancement of space technology, in recent years, inspires ambitious long-duration human space travel plan to space, Mars and asteroids [1]. To achieve such goals, it is suggested that the biotechnological solutions, based on engineered biochemistry of the cells, could be the key to the space technology challenges [2]. In one hand, small organisms, specially microbes, which are micron size replicating ‘reactors’, can survive in powder form (spore or lyophiazed) in various extreme condition including space [3], can be replicated fast when brings to appropriate environment, can be cultured in various types of reactors, and can produce a plethora of chemicals and materials [4–6]. On the other hand, engineered cells with synthetic genetic circuits produce highly controlled, human-designed functions [2,7,8]. Thus, it offers numerous projected applications for the near-future space technology, which spans from future space medicines, resource utilization, self-building habitats, space biomanufacturing, life support, programmable materials, and terraforming [2,9,10].

Microgravity is a unique and ubiquitous property of space, which is not experienced by the biochemical world on the earth. Microgravity influence biological processes at the molecular, cellular and organism level, including astronauts’ health and immunity [11–14]. Studies have been performed on multiple organisms to understand how they sense microgravity at the cellular and molecular level [11-13,15-20]. Several mechanisms have been proposed including the presence of gravireceptor [20], the effect of low shear stress [11, 21], altered transport phenomena due to microgravity induced low fluid shear [21], alteration of membrane fluidity [16] and the change in second messenger molecules [12]. However, no definite mechanistic details were known.

Biological solutions to space travel must consider microgravity as an important component. Thus, one of the fundamental challenges of space biotechnology is to create engineered biochemical systems to integrate microgravity as a physical signal within molecular and cellular processes. Engineered bacteria with synthetic genetic circuits have been shown to sense and integrate various environmental as well as intracellular chemical signals and compute complex human-designed functions through designed molecular interactions and reactions [22]. Apart from the chemical signals, cellular devices have been created to sense and integrate physical signals like temperature [23] and light [23, 24]. However, no molecular or biochemical microgravity-sensing device has been created. Here we created a synthetic biochemical device in living *Escherichia coli (E. coli)*, which can sense, integrate and respond to the microgravity. To our knowledge, this is the first biochemical or biological or molecular microgravity-sensing device and it shows a way to integrate microgravity as a physical signal with biochemical processes.

## RESULTS AND DISCUSSION

### Biochemical design principle of the microgravity sensing device

Though the microgravity brings prominent phenotypic changes in organisms including bacteria, the molecular genetic studies showed that the microgravity-induced alterations of expression of individual genes at both the transcript and the protein level were low [12, 18]. Here we target a small change in specific biomolecules in response to microgravity and process that signal through a molecular signal-processing device to get a bigger response and better control. Protein HfQ binds with a specific scaffold made with small regulatory RNAs (srRNA) and the resultant complex binds with target mRNAs to degrade it in bacteria including *E.coli* [25]. Using this principle, synthetic srRNA (SynsrRNA) based systems have been created to repress gene expression [25]. The intracellular quantity of HfQ within bacterial cells including *salmonella*, *pseudomonas*, and *E.coli* was reduced in microgravity [11, 18, 26].

To test, if the HfQ was down regulated in *E.coli* cell strain DH5αZ1 [27], chassis of our biochemical device, we ran the cells in High Aspect Ratio Vessels (HARV) in a rotating wall vessel (RWV) microgravity simulator, an important microgravity laboratory apparatus, designed by NASA [21]. When the reactors were placed vertically (Fig. 1a) the cells in a rotating wall vessel were subject to free-fall condition, which is a mimic of the microgravity. The same reactor in the horizontal position (Fig1b) gave the earth’s gravity (1G) control. *E.coli* DH5αZ1 in our experiment showed 50% less expression of HfQ mRNA expression in microgravity compare to the earth’s gravity control (Fig1c).

**Figure 1.**
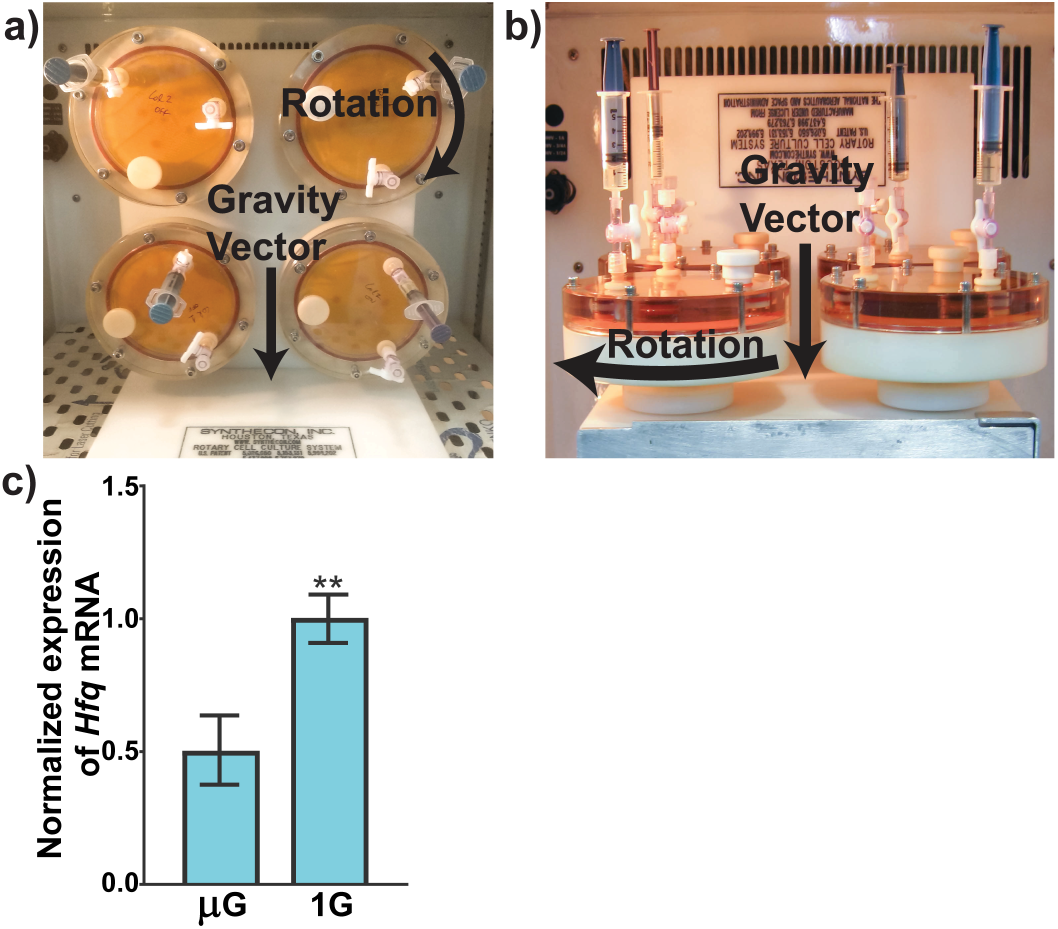
A) Working principle of microgravity simulating reactors. The volume of each of the RWV reactor is 50 mL. In vertical condition, they rotate in the shown direction. Thus, the cells within the reactors are always in free fall condition. B) For the Earth’s gravity control (1G) the reactors are placed in horizontal position. C) Relative HfQ mRNA expression level in microgravity (μG) and 1G. Three biological replicates were used for each of the gravity conditions.

We connected those two facts, as our design hypothesis. The decrease in HfQ protein in *E.coli* DH5αZ1 in microgravity would alter the repression capability of a designed SynsrRNA against a target protein expression compare to the earth’s normal gravity.

In that direction, we designed a SynsrRNA based molecular network (Fig. 2a), which may work as a microgravity sensing and responsive device in *E. coli*. Our designed SynsrRNA against enhanced green fluorescence protein (EGFP) mRNAwas around 130 bp long (FigS1) and its expression was under the control of an anhydrotetracycline (ATC) induced promoter pLtetO-1 [27]. The designed SynsrRNA consisted of an 87 bp MicC scaffold for HfQ binding [25] and a 40 bp sequence to bind the mRNA of the target enhanced green fluorescent protein (EGFP). The DNA sequence of the full SynsrRNA has been shown in supplementary figure 1 (FigS1). The EGFP is our machine-readable reporter protein, which was under the control of an Isopropyl β-d-1-thiogalactopyranoside (IPTG) regulated plLacO-1 promoter. The lacI and tetR proteins, which repress the plLacO-1 and pLtetO-1 promoter respectively, were produced constitutively in the *E.coli* cell strain DH5αZ1, the chassis of our device. Therefore, in presence of the IPTG only, the EGFP will produce and would give a high fluorescence signal. However, in the simultaneous presence of ATC and IPTG, the anti-SynsrRNA against EGFP mRNA would produce and reduce the EGFP expression and fluorescence signal.

**Figure 2.**
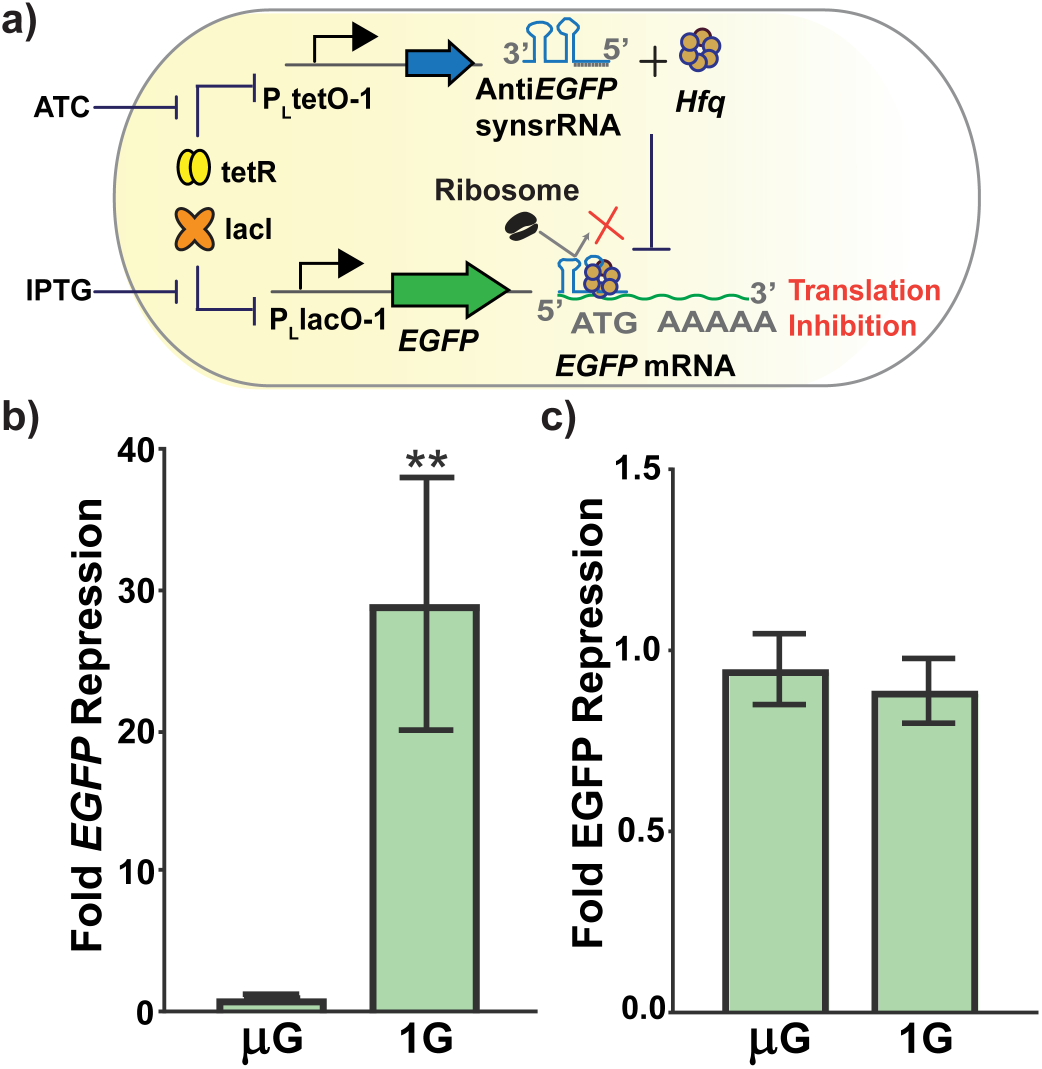
a) The design of the microgravity sensor circuit with EGFP. b) The experimental behavior of microgravity sensor. Each bar represent the EGFP expression folds change between IPTG only (absence) and IPTG + ATC (presence of anti-EGFP SynsrRNA) conditions. c) Control experiments.

The device was fabricated according to the design and incorporated in two different plasmid vectors, where plasmid L_eGFP_AC (FigS 2a) carries EGFP under IPTG inducible pLlacO-1 promoter in low copy (p15Ori) and plasmid T_antieGFP_synsrRNA_UA (Fig. S2b) carries SynsrRNA against EGFP mRNA under ATC inducible pLtetO-1 promoter in a very high copy (PUC Ori). Both the plasmids of our system were transformed into DH5αZ1 cells.

### The fabricated biochemical device works as a microgravity sensor

Next, the individual colonies of the engineered cells were subject to overnight growth with appropriate antibiotics at 37°C in a shaker incubator. Overnight cell culture from a single colony was divided into two groups. One group of cells was treated with both the inducers ATC and IPTG. The other group of cells was treated with IPTG only. The overnight culture was diluted 100 times and re-suspended in microgravity simulator reactors filled with fresh media with appropriate antibiotics and inducers. The cells in the simulators were grown for 24 hours at 37°C, collected, washed, re-suspended in PBS (pH:7.4) and the expression of EGFP was measured in a multimode micro-plate reader. We calculated (see method section) the fold changes in EGFP expression between IPTG only (express EGFP but not anti-EGFP SynsrRNA) and IPTG + ATC condition (express both EGFP mRNA and anti-EGFP SynsrRNA). A substantial differences in fold change (28.2) values between microgravity and normal gravity was observed (Fig. 2b). Next, to test, if the difference was stemmed from the microgravity induced changes in inherent expression of EGFP, instead of information processing through our device, we ran similar experiments with cells having EGFP under P_Llaco-1_ (FigS 2a) but without the plasmid carrying anti-EGFP SynsrRNA (FigS 2b), in two conditions (IPTG only and IPTG + ATC). We observed no significant differences (0.94 times, p value =0.4) between microgravity and normal gravity (Fig 2c). This suggested that our biochemical device sensed and integrated the microgravity signal and responded accordingly.

### Universality of the device design

As the SynsrRNA can be designed potentially against any gene (mRNA), our designed sensor should serve as a universal design for creating biochemical microgravity sensors. To test, we created a new biochemical device (Fig 3a) by replacing the target gene EGFP to TdTomato, an orange fluorescence protein, derived from DsRed, whose DNA sequence was completely different from EGFP and GFP derived fluorescent proteins [28]. Further, we replaced the anti-EGFP SynsrRNA to anti-TdTomato SynsrRNA (Fig S3) and the IPTG inducible PLacO-1 promoter to a constitutive lambda promoter P_R_. We performed the same microgravity experiments with this new system and found that it worked as a microgravity sensor too with a 3.77 times differences between 1G and microgravity (Fig 3b).

**Figure 3.**
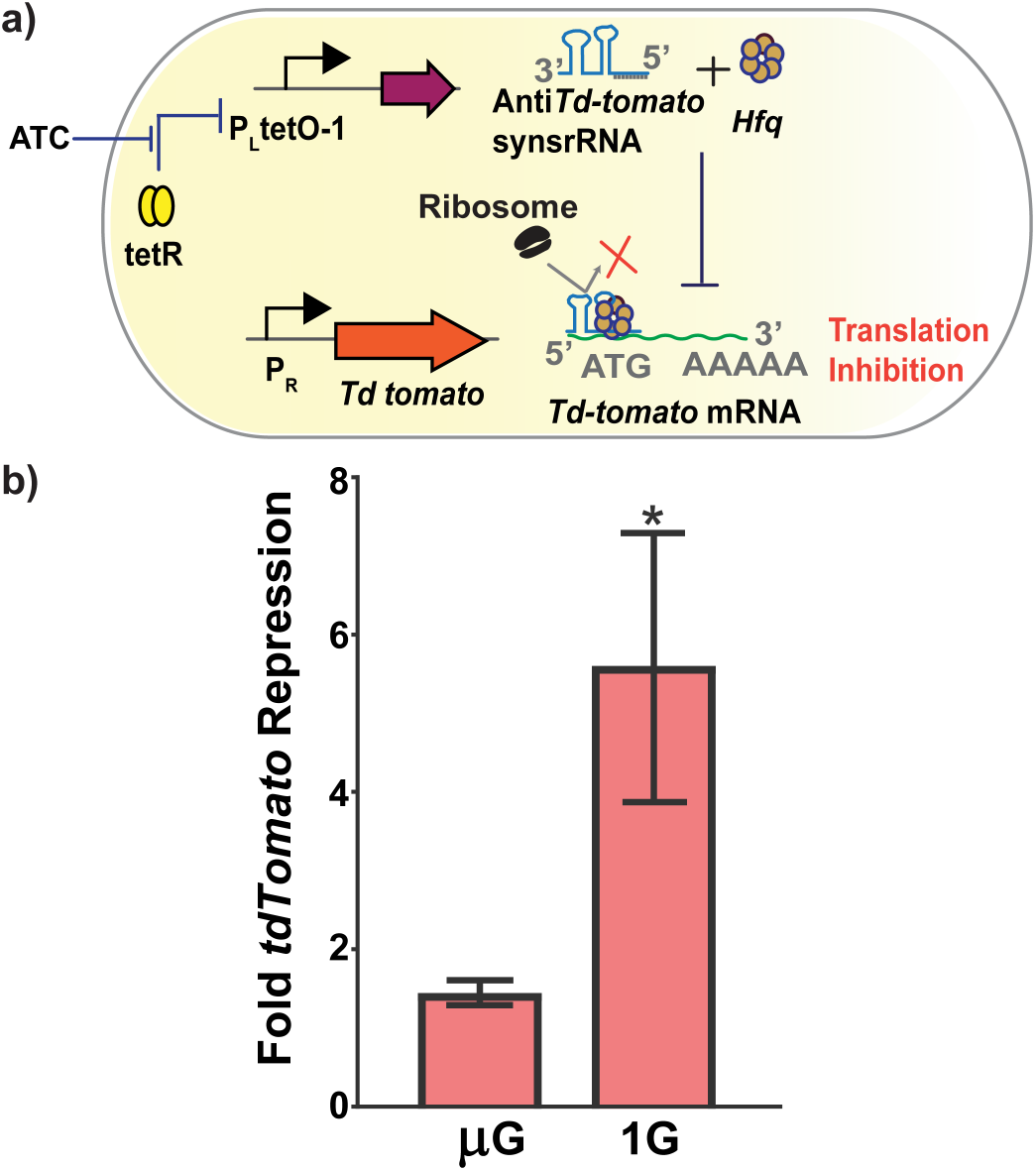
a) Gene circuit design of TdTomato based microgravity sensor. b) Experimental behavior of the TdTomato based microgravity sensor. Experimental characterization and calculation are same as in the EGFP based sensor.

### Application of the device to control native cellular properties with microgravity

As this biochemical device could integrate microgravity with it and potentially target any gene as a microgravity responsive gene, it may be applied to target inherent cellular biochemical process, which can be controlled in a microgravity responsive way. Therefore, we applied this device to control the cell division process with microgravity. FtsZ is a tubulin family protein, which is important for *E.coli* cell division. Its deregulation hinders the cell division process and shows elongated and filament-like shape [29]. Here we targeted native FtsZ by creating a microgravity responsive biochemical device (Fig S4a). The E.*coli* with the device expressing SynsrRNA against FtsZ showed that the deformed and elongated cell shapes in earth gravity got rescued in microgravity as shown in Fig. 4 and Fig S6a. However, the cell sizes in microgravity were still bigger than the control (FigS5 and FigS6b), where the engineered cells were grown in the absence of inducers in both gravity and microgravity conditions. All the individual fluorescence and DIC images are shown in FigS6.

**Figure 4.**
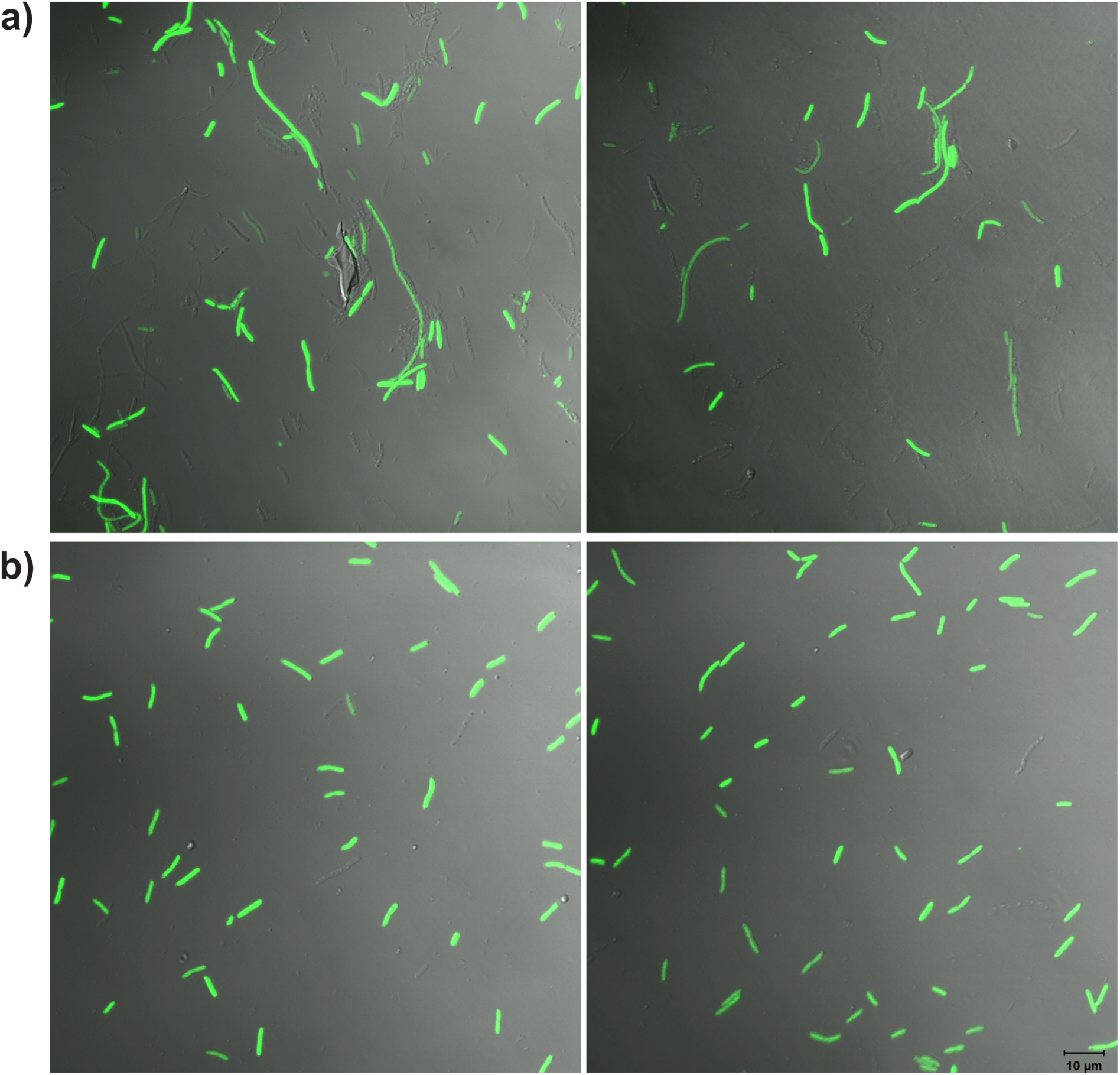
Merged fluorescence and DIC image of *E. coli* DH5αZ1 cells expressing SynsrRNA against FtsZ from microgravity responsive biochemical device were cultured in microgravity simulating reactor for 24 hours in presence of inducer at saturated concentration in A) Earth’s gravity condition (horizontal position), B) microgravity condition (vertical position).

## CONCLUSION

In this work, we created the first molecular or biochemical microgravity sensing and responsive device using an engineered genetic network in *E.coli*. We further demonstrated the universality of the device and its applications in integrating microgravity with inherent cellular biochemistry. Thus, it may open up a new direction in space bioengineering, where microgravity based cellular control would be important. One such application could be the development of ‘programmed space probiotics’ for astronauts’ against *salmonella* or *pseudomonas*, whose virulence has been enhanced in space microgravity and become a health hazard for astronauts in space [18]. Engineered probiotic bacteria may produce and secrete *salmonella* or *pseudomonas* killing peptides [30, 31] into astronauts’ gut only in the microgravity of space but not on the Earth.

Given that, the biochemical output from bacteria can be integrated with electro-chemical [32], mechanical [33] and optical systems [34], biochemical microgravity sensor will be useful to create material-cell hybrid robots [35] for various space medical technologies. Further, combining with an intracellular radiation sensor [23, 36] with a microgravity sensor in the same cell may give rise to a potential health hazards monitoring system in space.

Further, due to the universal nature of our design, the microgravity-sensing device can be integrated with any downstream synthetic and native cellular and metabolic pathways in *E.coli*. As HfQ proteins are downregulated in many bacterial species in microgravity, the microgravity responsive device can be built beyond *E.coli*. Taken together, our work may have a high significance in space biotechnology, space biochemical engineering, space chemistry, and gravitational biochemistry.

## MATERIALS AND METHODS

### Media, strains and growth conditions

We used DH5αZ1 strain of *E.coli* for our experiment as well as for cloning. The plasmid(s) were transformed in chemically competent *E. coli* DH5αZ1 cells and transformed cells were grown in LB-Agar, Miller (Difco, Beckton Dickinson) plates with appropriate antibiotics, followed by overnight liquid culture in Luria-Bertani, Miller broth (Difco, Beckton Dickinson) (LB) broth from single colonies at 37 °C with appropriate antibiotics. A single colony was picked and grown overnight in LB containing required antibiotic(s)(Ampicillin sodium salt, Himedia; Chloramphenicol, Himedia)at 37, 180 rpm. Next day the overnight culture was diluted 1/100 times in fresh LB containing required concentration of appropriate antibiotic(s) and inducer(s) (IPTG,Sigma Aldrich; ATC, Sigma Aldrich) and was transferred to High aspect ratio rotating vessels (HARVs; Synthecon, Houston, TX, USA) to mimic the microgravity environment (fig1a) at 37, 25 RPM for 24 hrs. We reoriented the previous setup in a horizontal position using a custom platform as a earth gravity control (fig1b). For cloning *E. coli* DH5αZ1 cells and transformed cells were grown in LB-Agar, Miller (Difco, Beckton Dickinson) plates with appropriate antibiotics, followed by overnight liquid culture in LB broth from single colonies at 37 °C with appropriate antibiotics.

### SynsrRNA and Plasmid construction

All the oligonucleotides used for the constructs are listed in table S2 and were synthesized from Integrated DNA Technologies, Singapore. The relevant plasmids maps are shown in fig. S2 and in table S1. *EGFP* gene, P_LtetO-1_ promoter,T1 terminator, ColE1 and p15A origin of replication (ori), ampicillin and chloramphenicol resistance genes were procured from plasmids: pOR-*EGFP*-12 and pOR-Luc-31 (a gift from Prof. David McMillen, University of Toronto, Toronto, Canada). The pUC origin of replication was amplified from pmCherry-N1 (Clontech) using pUC_Lp and pUC_Rp (Table S2). *tdTomato* gene was obtained from p*tdTomato* vector, clontech. P_R_ promoter along with RBS was constructed by overlap extension PCR using Pr_LpI and Pr_RpI. *EGFP* gene was fused to this promoter-RBS fragment by a overlap PCR with Pr_LpII and eGFP_RpII to construct full-length P_R_-*EGFP*. This P_R_-*EGFP* construct was flanked by *Xho*I and *Xba*I. The DNA encoding for small synthetic RNA against *EGFP* was chemically synthesized (Invitrogen GeneArt Gene Synthesis service, Thermo Fischer), the same was used as a template to construct synsrRNA against *tdTomato* using primer antitom_Lp and antitom_Rp. The SysrRNA against *Ftsz* was constructed by overlap pcr of primer antiftsz_Lp and antiftsz_Rp. All SynsrRNA were cloned downstream of P_LtetO-I_ promoter using EcoRI and Xba-I into plasmid harbouring pUC origin and ampicillin resistance. The target genes *EGFP* and *tdTomato* gene were introduced in a plasmid with p15a origin and chrolamphenicol resistance gene. The plasmid containing the synsrRNA e.g anti*EGFP*synsrRNA and the corresponding target gene e.g. *EGFP* were co-transformed in DH5αZ1. To aid visualization a plasmid P_*EGFP*_AC, where *EGFP* is constitutively expressed under P_R_ promoter, was used in case of ftsz silencing experiment. All PCR reactions were carried out by KOD Hot Start DNA polymerase (Merck Millipore). PCR and Gel purification were done using Qiaquick nucleotide removal kit (qiagen) and Oiaquick Gel purification kit (qiagen), respectively. All the plasmids were extracted using qiaquick miniprep kit (qiagen) and sequenced from Eurofins Genomics India Pvt. Ltd., Bangalore, India.

### Fluorescence Measurements and data analysis

At least 3 biological replicates were used for comparing fold repression between microgravity and earth’s gravity. Cell fluorescence and absorbance at 600 nm were measured after 24 hrs of culture in microgravity simulator. The cells were harvested and washed twice in PBS (pH  7.4). Then the cells were diluted in PBS (pH  7.4) to reach around OD_600_ 0.8, loaded onto 96-well multiwell plate (black, Greiner Bio-One) for measurement using Synergy HTX Multi-Mode reader (Biotek Instruments, USA). For measuring fluorescence of EGFP, the cells were excited by a white light source that had been passed through an excitation filter 485/20  nm and emission was collected by 516/20  nm bandpass filter. Similarly for fluorescence measurements of tdTomato an excitation filter of 540/35  nm and an emission filter of 590/20  nm was used. The OD_600_ was also measured in the same instrument.

The raw fluorescence measurements were normalized using equation 1,

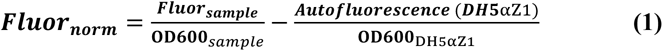

where ***Fluor_norm_*** is the normalized fluorescence, ***Fluor_sample_*** is the raw fluorescence of the sample, **OD600**_*sample*_ is the absorbance of the sample at 600nm, ***Autofluorescence***_***DH*5**αZ1_ is the auto-fluorescence of DH5αZ1 strain, and **OD600**_***DH*5**αZ1_ is the absorbance of DH5αZ1 strain at 600nm.

The *average fold repression* of EGFP and Td-tomato expression between the IPTG only and IPTG + ATC condition was calculated as a ratio between average normalized fluorescence over multiple biological replicates in uninduced condition and that of induced condition.

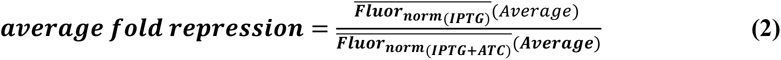

The 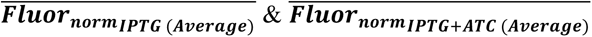 are the average *EGFP or tdTomato* normalized fluorescences over multiple biological replicates for IPTG only and IPTG + ATC condition, respectively. Unpaired t-test (double tail) was used to measure the extent of statistical significance. All data were analyzed and plotted using GraphPad Prism 8.

### RNA purification and cDNA sysnthesis

The Bacterial cells were treated with RNAprotect bacteria reagent (qiagen) and kept at −80°C until RNA extraction. The bacterial cells were lysed using lysozyme (sigma) and proteinase-k (qiagen) treatment and the RNA was extracted using RNAeasy mini kit (qiagen), following manufacturer’s instruction. To lower the chance of genomic DNA contamination On-column DNASE digestion step was implemented using RNase free DNAse (qiagen). The extracted RNA was denatured using formamide and ran in 1.2% agarose gel for checking its quality. The concentration of the RNA was measured using UV-Vis absorbance function of Biotek synergy HTX plate reader. One microgram of RNA was used for cDNA synthesis using Qiagen quantitect reverse transcription kit according to the manufacturer’s instruction.

### qPCR and Data analysis

All oligos used for qpcr are listed in and were synthesized from Integrated DNA Technologies, Singapore. The qPCR was carried out using qiagen quantitect SYBR green PCR kit, according to manufacturer’s instruction in applied biosystems step one plus real time PCR system. Atleast three biological replicates and three technical replicates were used for qPCR. The qPCR was performed for hfq and 16s rRNA as a endogenous reference gene as used previously elsewhere[34]. A no template control showed the absence of detectable transcipts for both primer set. The relative expression values using standard ΔΔCt method.

### Confocal microscopy

After 24 hrs the bacterial cells were harvested at 4000 rpm and washed thrice in PBS. The cells were finally diluted in PBS enough to prevent crowding. To visualize, ten microliters cell suspension was put onto an one percent thin agarose pad (SeaKem LE agarose) and was covered with a cover slip. The DIC and fluorescence imaging was performed in zeiss in LSM 710 confocal microscope, where the pinhole was kept partially open and the images were captured using zen 2008 software.

## ACKNOWLEDGEMENTS

This work was supported by IBOP Project (Department of Atomic Energy, Govt. of India), and Ramanujan Felowship (DST), Govt. of India

## Supporting Information

**Fig S1.**
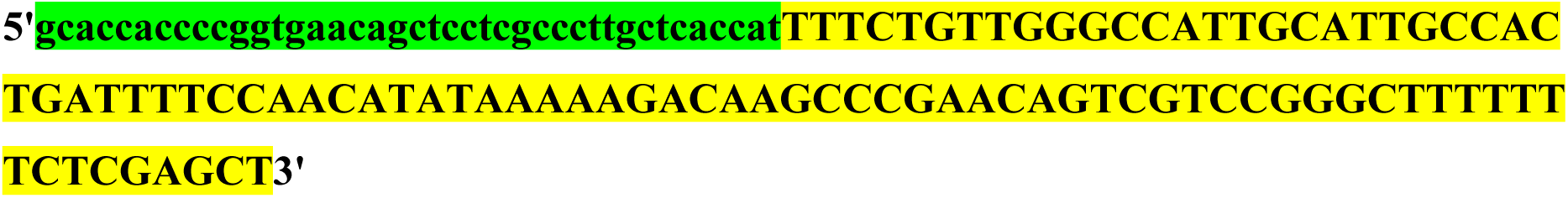
Anti-*EGFP* synthetic small regulatory RNA sequence. The sequence highlighted in green is the target (EGFP mRNA) binding region and the sequence highlighted in yellow is the MicC scafold region, which helps to bind Hfq protein.

**Fig S2.**
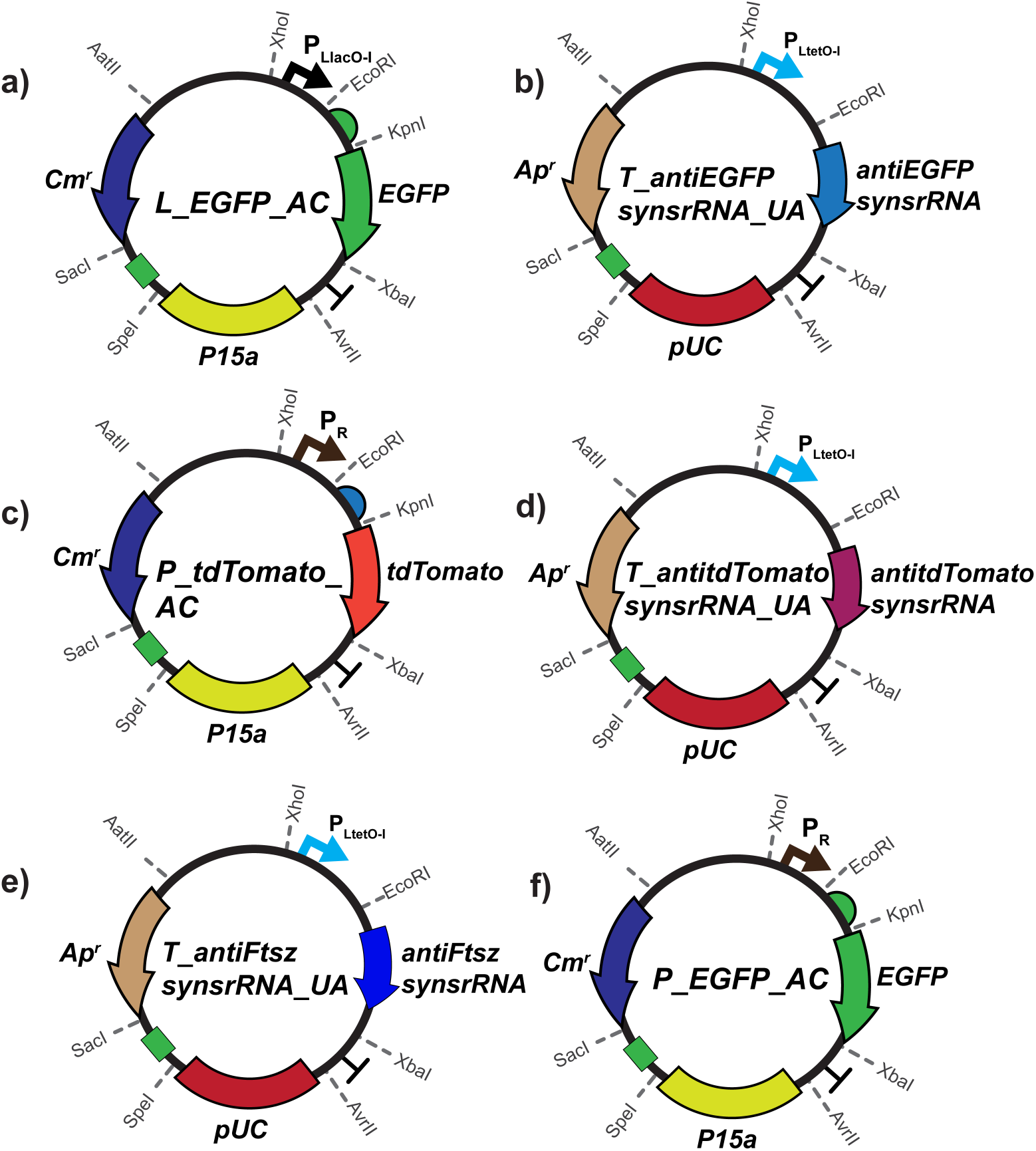
Plasmid maps for Important plasmids used in this study.

**Fig S3.**
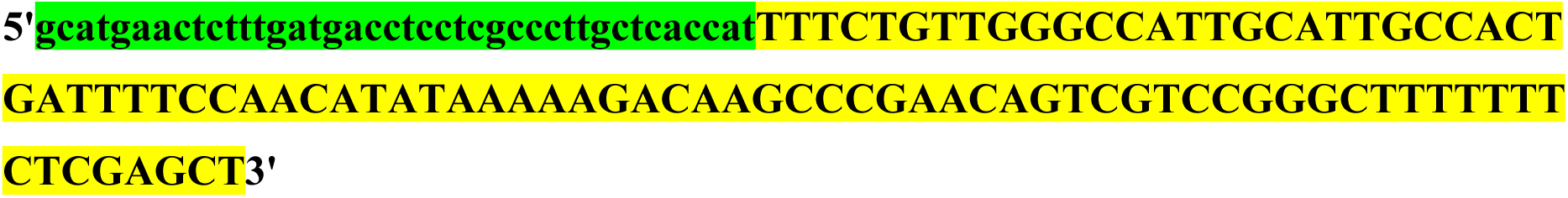
Anti-*tdTomato* synthetic small regulatory RNA design. The sequence highlighted in green is the target (tdTomato mRNA) binding region and the MicC scaffold is highlighted in yellow.

**Fig S4.**
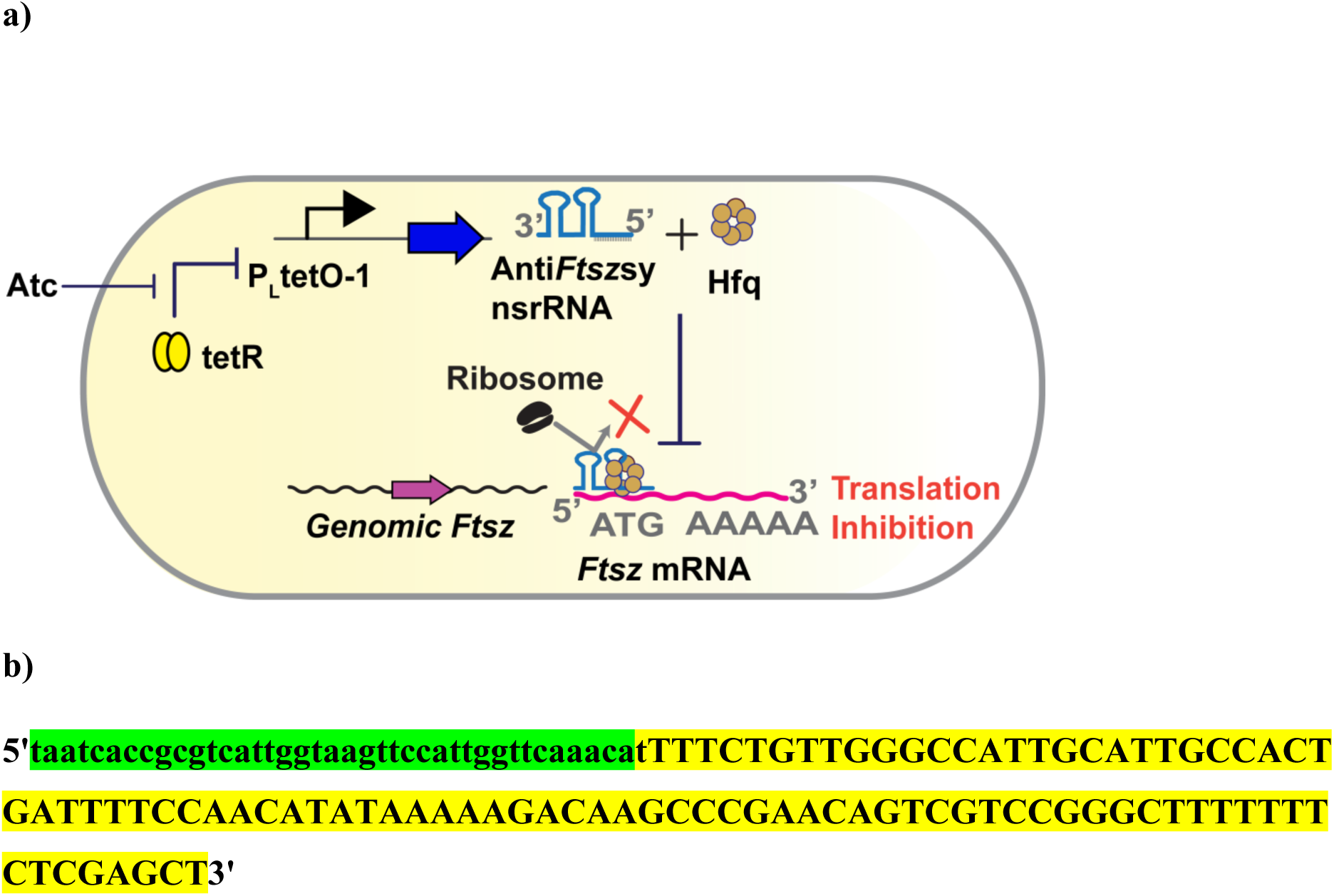
**a)** SynsrRNA based microgravity sensor for integrating microgravity with native FtzS protein in Dh5αZ1 *E.coli.* **b)** Anti*Ftsz* synthetic small regulatory RNA design. The sequence highlighted in green is the target binding region and the MicC scaffold is highlighted in yellow.

**Fig S5.**
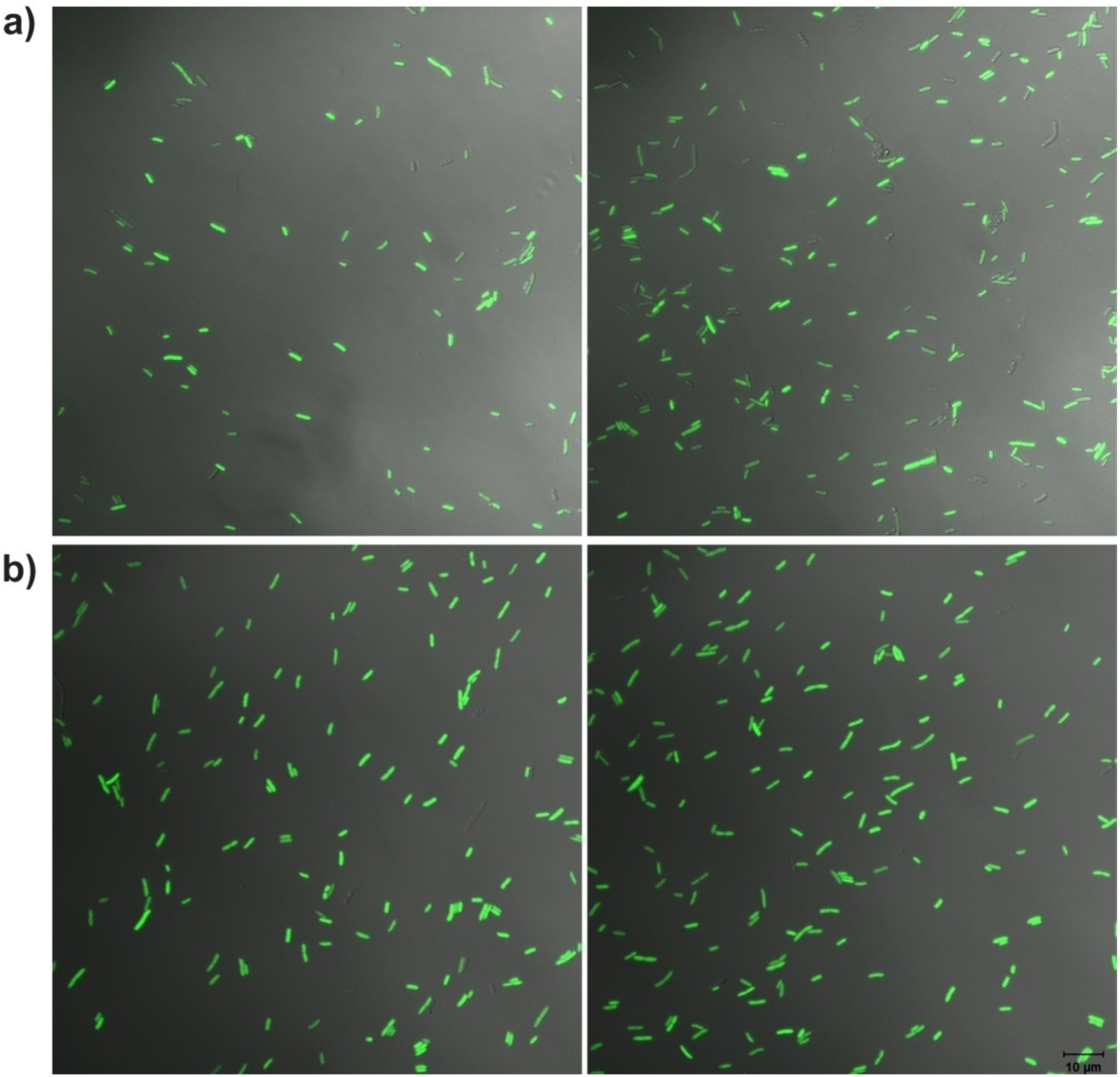
**Control experiments for Figure 2 in main text:** Merged images of Dh5αZ1 cell co-transformed with anti-*Ftsz* synsr, expressing SynsrRNA against genomic FtsZ under Atc inducible PltetO-1 promoter and P_EGFP_AC, expressing EGFP constitututvely from P_R_ promoter in absence of inducer, Atc at a) Earth’s gravity (RWV at horizontal position) b) microgravity (RWV at vertical position)

**Fig S6.**
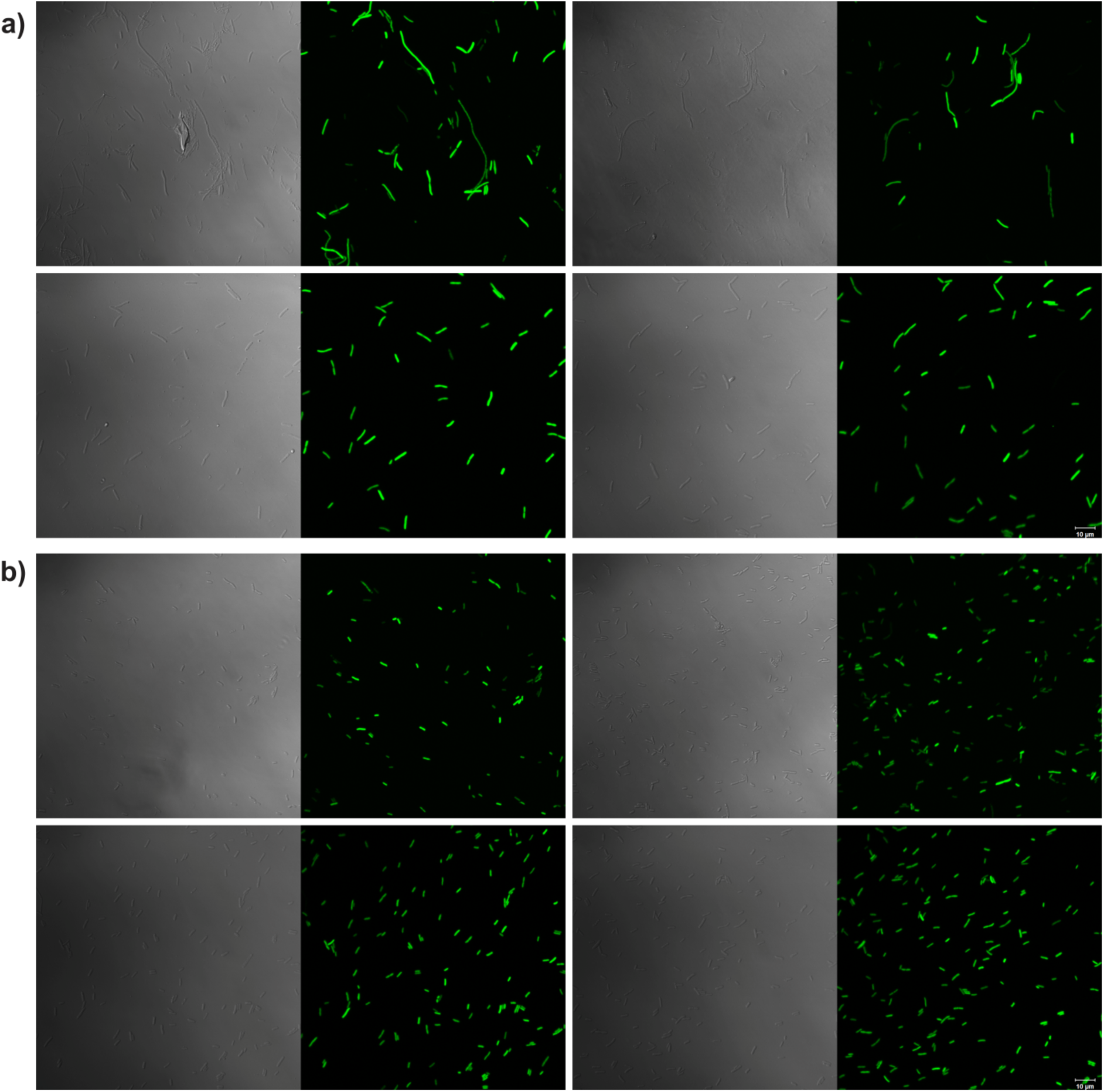
DIC and Flouresence image of the merged images a) in Fig 2 in main text and b) in FigS5 (control experiments).

**Supplementary table 1 :**
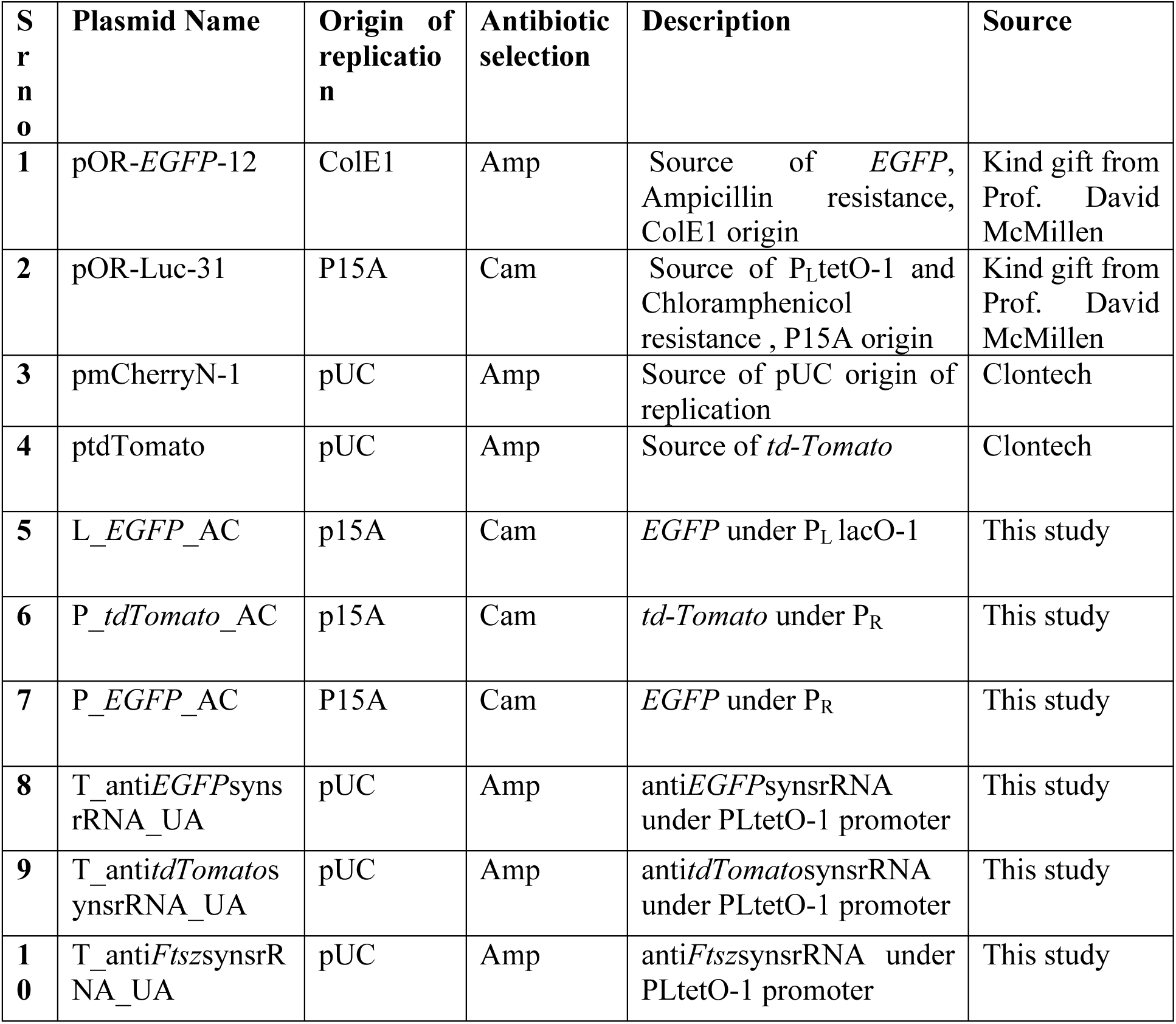
List of all the plasmids used in this study.

**Supplementary table 2 :**
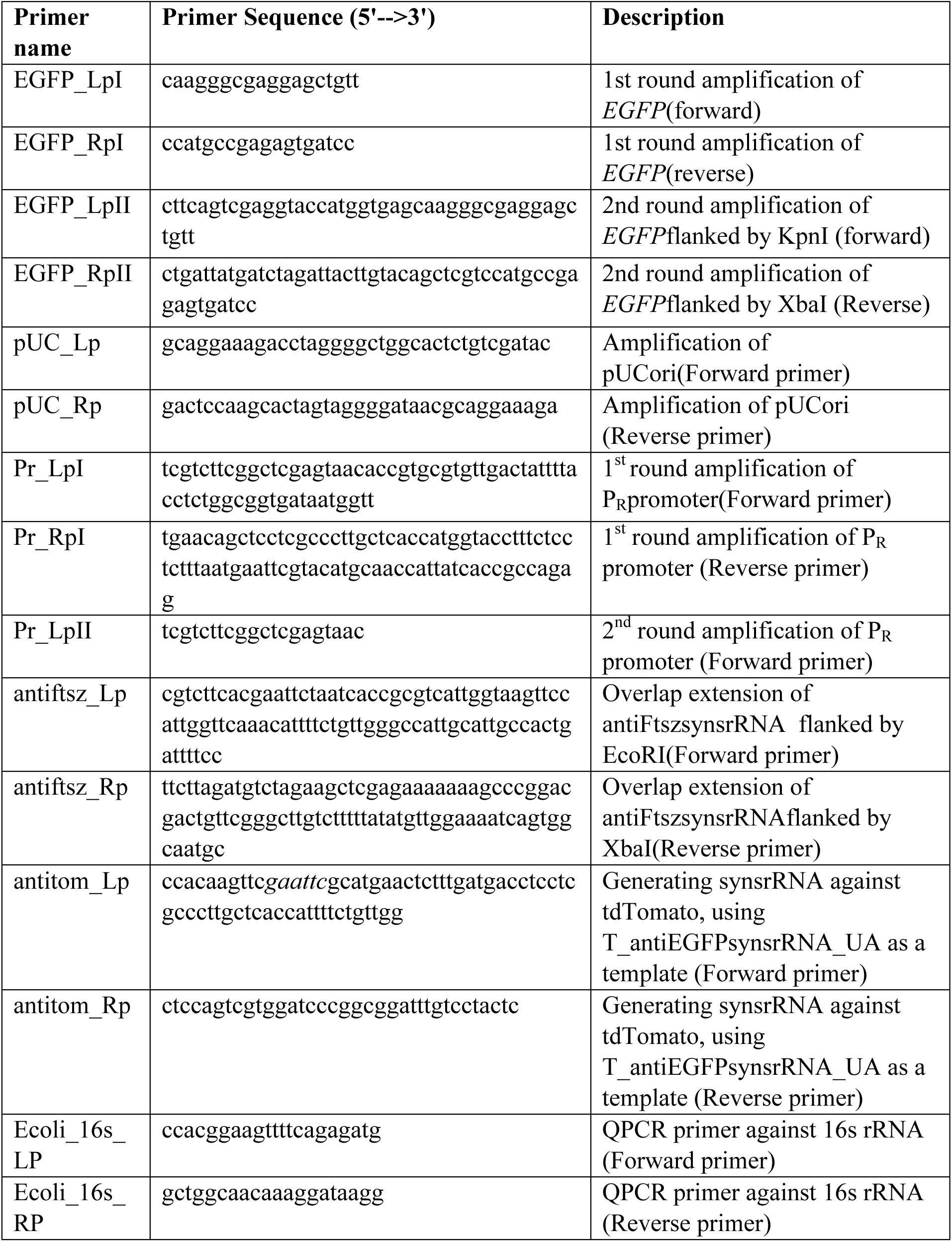
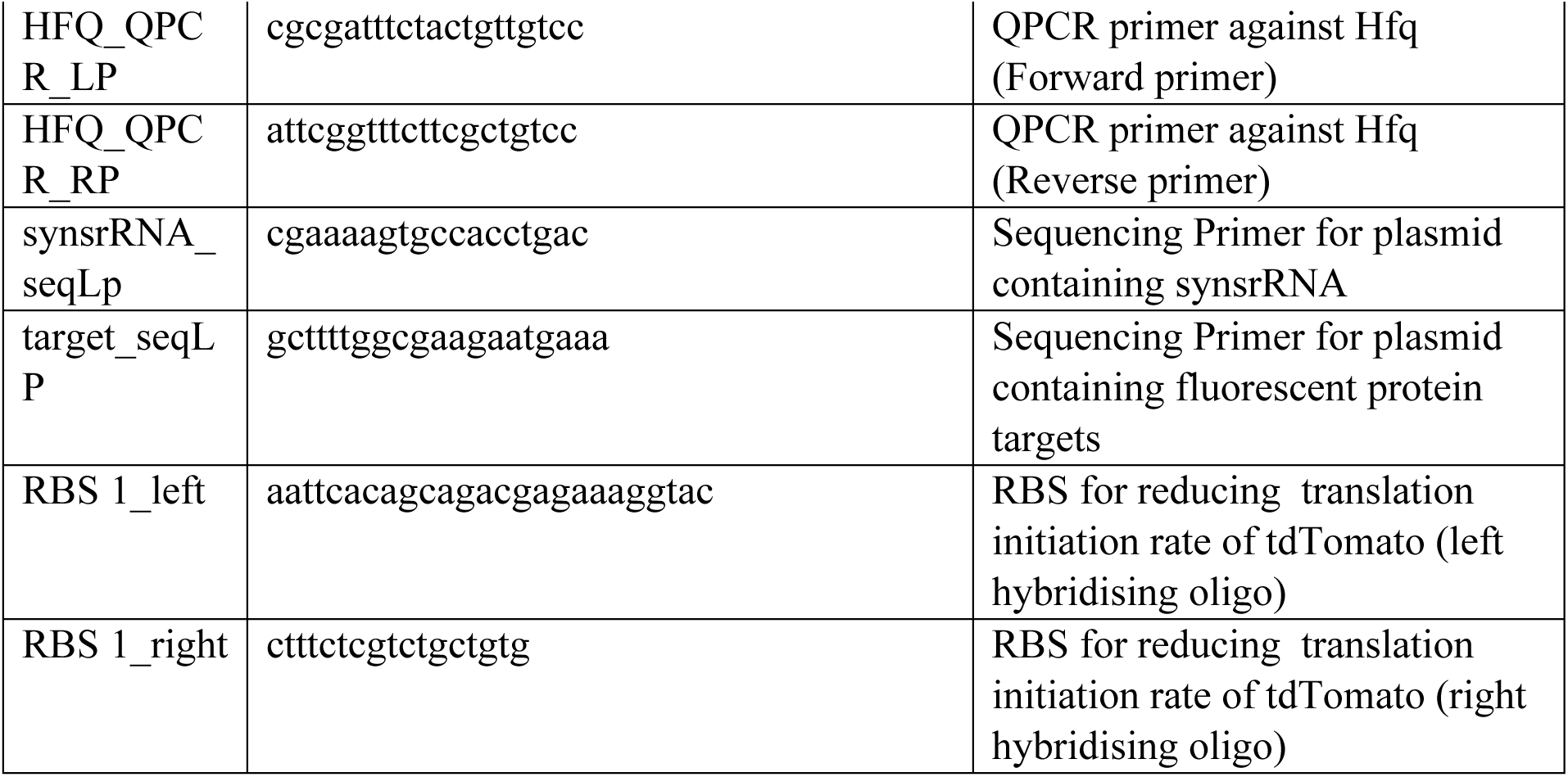
List of all the primers used in this study.

